# Topographically organized representation of space and context in the medial prefrontal cortex

**DOI:** 10.1101/2021.06.04.447085

**Authors:** Jonas-Frederic Sauer, Shani Folschweiller, Marlene Bartos

## Abstract

Spatial tuning of neocortical pyramidal cells has been observed in diverse cortical regions and is thought to rely primarily on input from the hippocampal formation. Despite the well-studied hippocampal place code, many properties of the neocortical spatial tuning system are still insufficiently understood. In particular, it has remained unclear how the topography of direct anatomical connections from hippocampus to neocortex affects spatial tuning depth, and whether the dynamics of spatial coding in the hippocampal output region CA1 such as remapping in novel environments is transmitted to the neocortex. Using mice navigating through virtual environments we addressed these questions in the mouse medial prefrontal cortex, which receives direct input from the hippocampus. We find a rapidly emerging prefrontal representation of space in the absence of task rules, which discriminates familiar from novel environments and is reinstated upon re-exposure to the same familiar environment. Topographical analysis revealed a dorso-ventral gradient in the representation of the own position, which runs opposite to the innervation density of hippocampal inputs. Jointly, these results reveal a dynamically emerging and topographically organized prefrontal place code during spontaneous locomotion.

**Significance Statement:** The neocortex is composed of areas with specialized function (e.g. sensory vs. associational). Despite this functional diversity, emerging evidences suggest that the encoding of space might be a universal feature of cortical circuits. Here, we identified a gradient of spatial tuning depth along the dorso-ventral axis. A complex topography of spatial tuning properties might support a division of labor among medial prefrontal cortical subnetworks to support local circuit computation relevant for the execution of context-dependent behavioral outcomes.

## Introduction

Encoding of spatial information is classically assigned to the hippocampal-entorhinal system (1) (2) (3). However, with recent reports of spatially tuned neurons in the posterior parietal (4), retrosplenial (5), visual (6) (7), somatosensory (8) and orbitofrontal (9), as well as over large areas of the dorsal cortex including sensory and higher-order areas (10), it is becoming evident that spatially modulated activity is a more broadly appearing phenomenon than initially assumed, suggesting that spatially tuned input might be relevant for local information processing.

The medial prefrontal cortex (mPFC) is critical for cognitive functions such as the planning and execution of complex behaviors (11). Many cognitive tasks involving the mPFC require the simultaneous awareness of spatial variables including one’s own current position within an environment, the recognition of novelty/familiarity of the surrounding features, and context discrimination (12) (13). In line with this, prefrontal neurons of mice display spatially tuned activities during cognitive tasks (14) (15) (16) (17) (18). However, compared to the extensively studied properties of spatial coding in the hippocampal-entorhinal system, several important questions about spatial tuning in the mPFC remain open. First, the majority of previous work reported spatial tuning in cognitive tasks in which the animals had learned reward locations (14) (15) (16) (17) (18). In contrast, little evidence of spatial tuning was found in spontaneously exploring rodents (17) (19). These findings suggest that for spatially tuned activities to emerge, the mPFC network might need to be engaged in active rule learning. Second, neurons in hippocampal CA1 display global remapping (i.e., a change in the location of the firing field of the neuron) when the animal enters a novel environment (20). The mPFC receives direct input from CA1 (21) (22), but it is unclear whether mPFC neurons inherit context-dependent remapping from CA1 afferents. Finally, afferent input from the hippocampal formation targets predominantly ventral prefrontal regions (23) (24) (25) (26), raising the question whether spatial tuning depth in the mPFC increases along the dorso-ventral axis as one might predict based on the topography of monosynaptic hippocampal inputs.

Here, we addressed these questions by assessing spatial tuning in the mPFC along the dorso-ventral axis while mice were spontaneously navigating through virtual environments. We find that prefrontal pyramidal cells are spatially tuned when animals explore familiar or novel environments in the absence of rewards or rule learning. Prefrontal neurons remap when the animal is introduced to a novel arena, while the original place code is partially reinstated upon return to the familiar environment. Finally, we find a topographic gradient in the spatial modulation depth and consistency in spatial tuning of prefrontal pyramidal cells along the dorso-ventral axis, which runs in the opposite direction as the density of hippocampal inputs. Our data, thus, reveal a rapidly emerging and topographically organized prefrontal place code during spontaneous locomotion.

## Results

### Prefrontal neurons are spatially tuned during spontaneous unrewarded navigation

We trained head-fixed mice to run consecutive laps on a virtual circular track (1.5 or 2 m long; **Fig. 1**). Since we observed similar spatial tuning for recordings on the 1.5 and 2 m tracks, the data were pooled (**SI Appendix, Fig. S1**). The track was composed of linear segments with distinctive wall patterns and additional cues located outside the track (**Fig. 1A**). We chose this experimental setting to constrain the assessment of spatial tuning in one dimension (i.e., 1-D position on the track) and to avoid complex behaviours such as grooming or rearing, which are frequently observed in freely moving mice and modulate the activity of prefrontal neurons (27). The animals moved along the track in a self-induced and self-paced manner without rewards. After familiarization to the track for several days, we recorded single units with linear silicon probes acutely lowered into the mPFC (**Fig. 1B**).

**Figure 1:**
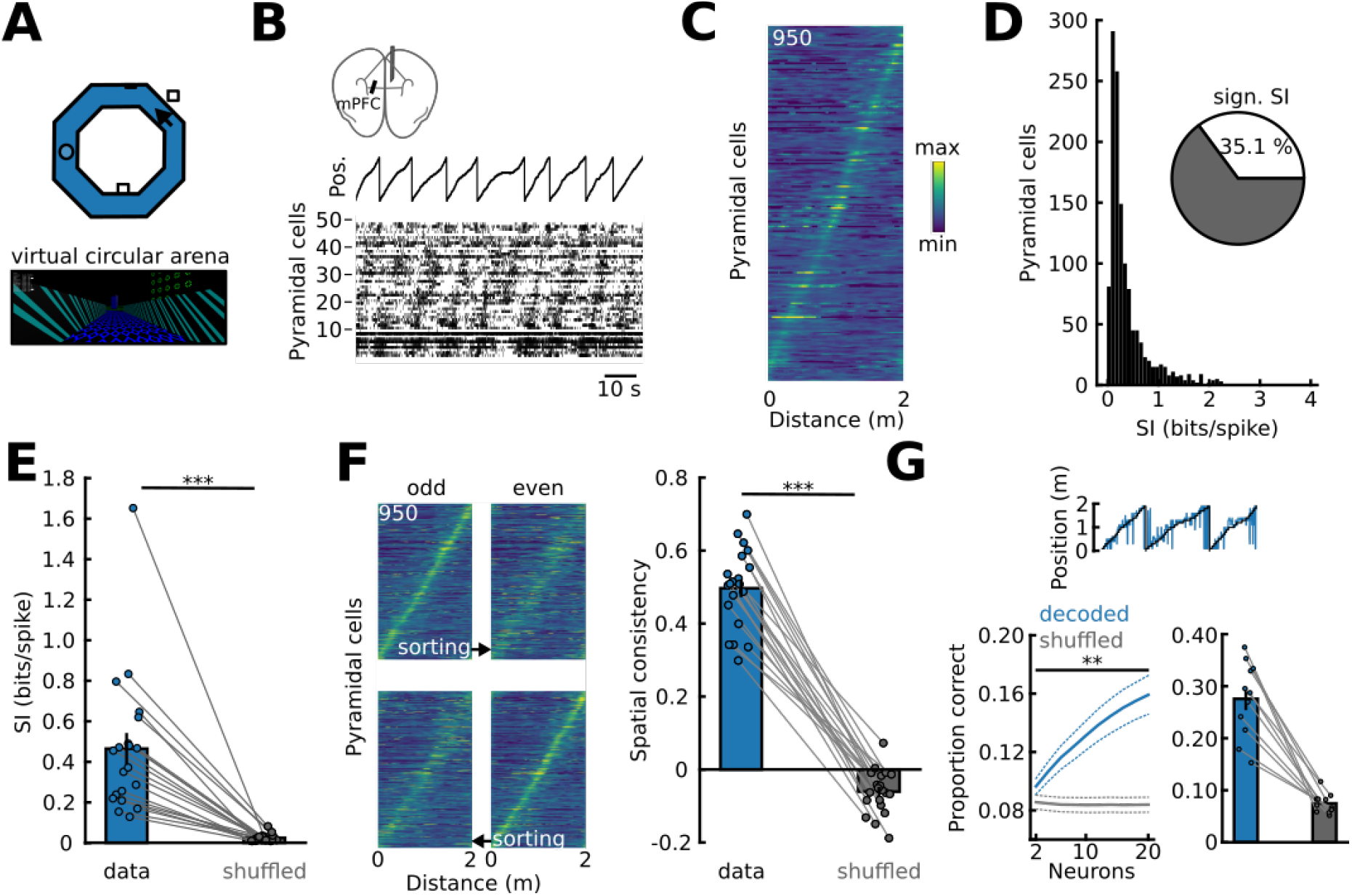
Spatially tuned activity of prefrontal pyramidal neurons during self-paced, unrewarded motion. (A) Experimental protocol for head-fixed recordings during navigation in the virtual reality. Arrow on the track indicates the position shown at the bottom. (B) Single-unit recording during movement on the track. Black line indicates position on the track. Each row of the raster plot denotes one unit sorted for the location of peak activity. (C) Spatially binned activity of pyramidal cells on a 2 m familiar track (*N*=15 mice). See **SI Appendix Fig. S1** for similar results for 5 additional mice running on a 1.5 m track. (D) Histogram of spatial information content (SI) of individual pyramidal cells. Inset, circular plot shows that 35.1% of recorded cells expressed a significant SI (**Methods**). (E) Pyramidal cells within individual mice show significant mean spatial tuning compared to shuffled controls. *P*=1.3*10^−5^, *N*=20 mice, paired *t-*test. (F) Significant consistency of spatial tuning: Left, tuning function during alternating runs sorted for odd (top) or even runs (bottom). Right, spatial run-by-run consistency (i.e., correlation of spatial tuning functions between odd and even runs) was significantly different from shuffled control data. *P=6*.*5*\*10^−14^, *N*=20 mice, paired *t-*test. (G) Position could be predicted with a linear support vector classifier based on spike rates of randomly selected pyramidal neurons. *P=*9.5*10^−6^-0.001, paired *t-*tests with Bonferroni correction for 10 comparisons, *N*=17 mice. Right, decoding with maximal cell number in sessions with >30 pyramidal neurons (average number of used neurons: 56 ± 4). *P=*1.6*10^−6^, *N=*11 mice, paired *t-*test. Points in (E), (F) and (G) show averages for individual mice. Circles connected by lines were obtained within one animal. Continuous and dotted lines in (G) represent animal means ± sem.

In total, we recorded from 2329 active prefrontal neurons (*N*=20 mice). Based on spike waveform kinetics, we grouped the cells in putative pyramidal (*N*=2005) and in GABAergic interneurons (*N*=324, **SI Appendix, Fig. S2**). To assess spatial representation, we quantified spatial information (SI) from spatially binned firing rates (28). SI of putative interneurons was lower than that of pyramidal cells (**SI Appendix, Fig. S2**). We therefore focused on pyramidal cells for the remainder of the study.

Analysis of SI revealed a broad distribution for the pyramidal cell population, with many units displaying spatially confined activity (all pyramidal neurons that discharged at least at 1 Hz during movement were considered for analysis, *N=*1272 neurons; **Fig. 1B-D**). Comparison against 1000 shuffled spike trains revealed that 35.1 % of all recorded prefrontal pyramidal neurons displayed significant SI (**Fig. 1D, SI Appendix, Table S1**). Consistent with previous reports in spatial memory tasks (14), spatial tuning of prefrontal pyramidal cells was lower than in the hippocampus (*P=*0.001; **SI Appendix, Fig. S3**). However, the average SI of pyramidal cells was significantly different from a surrogate dataset created by locally shifting the spike trains of each unit in time on the level of individual mice, indicating above-chance spatial representation by prefrontal pyramidal neuron populations during spontaneous locomotion (*P*=1.3*10^−5^, paired *t-*test, *N=*20 mice; **Fig. 1E**).

We conducted additional analyses to assess the validity of our spatial tuning measure and the robustness of the observed spatial tuning during spontaneous locomotion. First, significant SI of the population was consistently observed against different shuffled distributions and for different spatial binning methods (**SI Appendix, Fig. S4**). Second, movement speed or acceleration were unlikely to have impacted spatial tuning since above-chance SI was also evident when only neurons without significant speed modulation were considered (∼40% of pyramidal neurons were speed-modulated; **SI Appendix, Fig. S5**). Third, we found significant consistency of the spatial tuning when correlating the spatial tuning functions for odd and even runs within a session versus the shuffled spike trains (mean correlation: 0.49 ± 0.02, *N=*20 mice, *P*=6.5*10^−14^ versus shuffled spike trains, paired *t-*test; **Fig. 1F**). Moreover, significant consistency was observed when we assessed consistency based on runs of the first versus the second half of the recording session (**SI Appendix, Fig. S6**). Finally, decoding with an increasing number of randomly chosen pyramidal cells allowed the progressively increasing prediction of the animal’s position compared to shuffled data (**Fig. 1G**). Thus, the activity of prefrontal pyramidal neurons contains information about the animal’s position during spontaneous self-paced motion.

### Prefrontal neurons remap in a novel context but regain their tuning in the familiar environment

To assess the stability of prefrontal spatial tuning across contexts we exposed mice to a sequence of familiar (fam), novel (nov), and again the familiar track (fam’) within one recording session (**Fig. 2A**,**B**). The novel track differed from the familiar environment in size (track length 3 m) and visual features (colors, wall patterns, external cues, **Fig. 2A,B**). We previously demonstrated that mice are capable to recognize such novel virtual tracks (20). Analysis of waveform parameters indicated stable recording conditions throughout the exposure to the three environments (**SI Appendix, Fig. S7**). The neurons displayed comparable average firing rates during running in all three experimental conditions (**Fig. 2C**).

**Figure 2:**
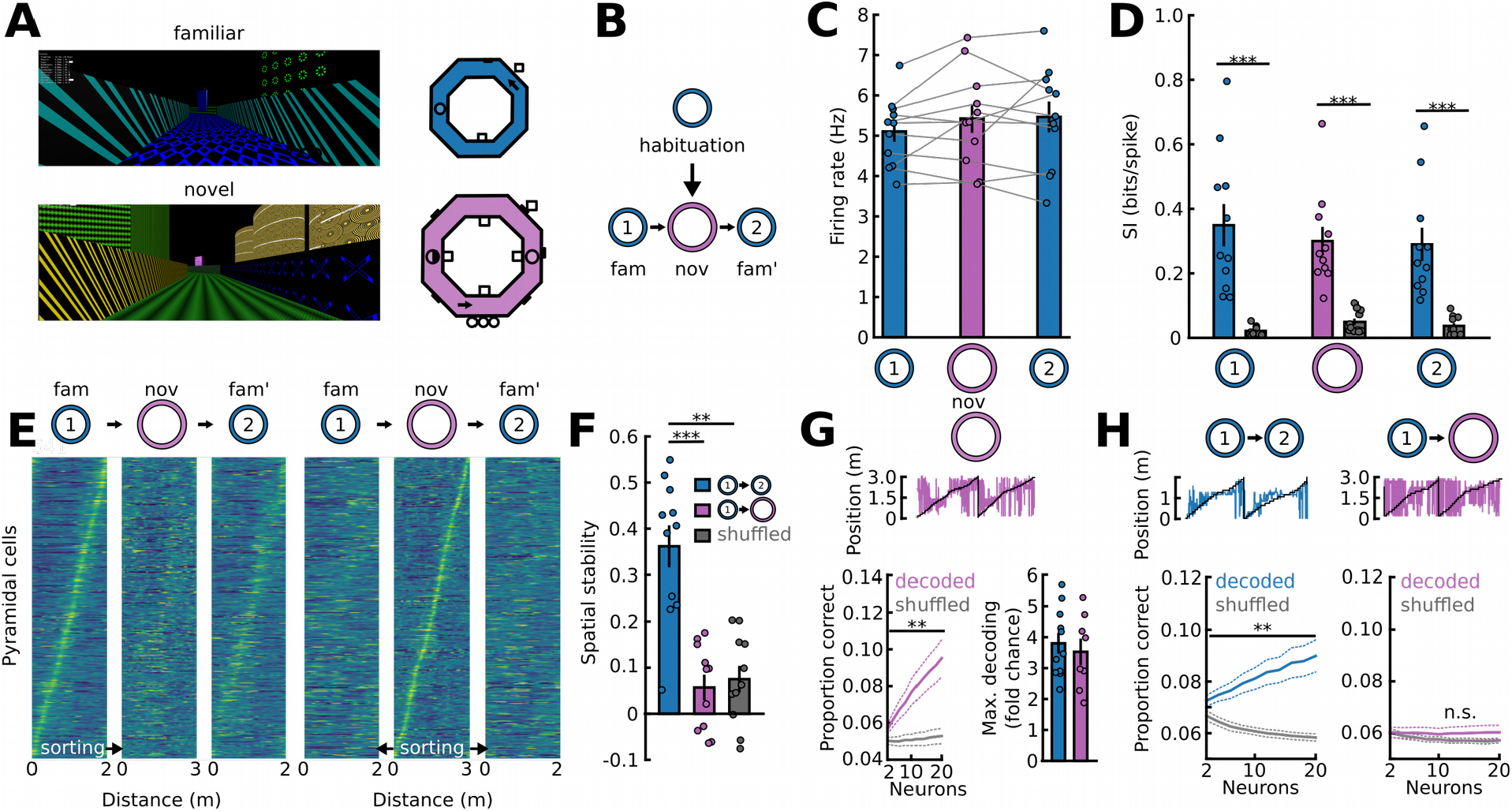
Prefrontal spatial tuning remaps in a novel environment. (A) The animals were sequentially exposed to a familiar (blue, fam) and novel track (purple, nov). Right: Schematic of the two tracks. Arrows indicate position of the views shown on the left. (B) Schematic of the recording paradigm. (C) Pyramidal cells within individual mice displayed comparable mean firing rates during all three conditions. *P=*0.711, one-way ANOVA. (D) Prefrontal units showed significant SI in all three conditions. Fam: *P*=0.0004, nov: *P*=5.7*10^−5^, fam’: *P=*0.0003, paired *t-*tests, *N=*11 mice. (E) Tuning maps of pyramidal cells during the fam-nov-fam’ sequence sorted for their peak location in the fam (left) or nov (right) environment. (F) Spatial stability of the tuning function was significantly higher for fam-fam’ than for fam-nov pairwise comparisons or shuffled controls. *P=*2.4*10^−5^ versus nov and *P=*0.003 versus shuffled, *N=*11 mice, paired *t-*tests with Bonferroni correction for three comparisons. (G) Decoding position in nov. Top: Example of decoded trajectory. Bottom left: The animal’s position could be significantly decoded from pyramidal cell spike trains during exploration of nov. *P=*0.003-0.013, *N=*10 mice, paired *t-*tests with Bonferroni correction for 10 comparisons. Bottom right: Similar maximal decoding accuracy using all neurons in fam (56 ± 4 cells) and nov (57 ± 7 cells, *P=*0.295 for the comparison of neurons used for decoding, Mann-Whitney U test). Data are expressed as fold chance level estimated from shuffled data. *P=*0.614, *N=*11 and 8 mice (H) Training the decoding model on fam allowed the significant prediction of the animal’s position in fam’ (left, *P=*0.0002-0.002) but not in nov (right, *P=*0.201-0.684), paired *t*-tests with Bonferroni correction for 10 comparisons. Circles in (C), (D), (F) and (G) show average values for individual mice. Circles combined with lines represent data from individual mice. Continuous and dotted lines in (G) and (H) show animal means ± sem.

While pyramidal neurons showed significant SI compared to shuffled datasets in all three conditions on the level of individual mice (**Fig. 2D**), correlation of the neuron’s spatial tuning functions between fam and nov was low (mean correlation 0.057 ± 0.028, *N*=11 mice; **Fig. 2E,F**). These data suggest that exposure to a novel environment leads to a rearrangement of spatial representation in the mPFC (i.e., to remapping). Indeed, when sorted for peak activity, the firing of prefrontal neurons tiled the entire track in the novel environment (**Fig. 2E**). Furthermore, decoding analysis allowed the significant prediction of the animal’s position in nov with similar accuracy as in fam, confirming that cells rapidly formed a new spatial representation in the novel environment (**Fig. 2G**).

We next asked to what extent the spatial tuning reverts to the original representation upon re-exposure to the familiar environment after exploration of nov. Compared to the correlations to nov or to shuffled data, we found a significantly higher correlation to the original spatial pattern observed in fam (mean correlation: 0.362 ± 0.045; *P=*2.4*10^−5^ versus nov and *P=*0.003 versus shuffled, *N=*11 mice, paired *t-*tests with Bonferroni correction for three comparisons; **Fig. 2F**), indicating that the spatial representation of fam is partially reinstated on the level of individual mice. To corroborate this finding, we used a decoder trained on the activity of a randomly chosen subset of neurons in fam to predict the position on the track in fam’ or nov. This analysis revealed significant decoding compared to shuffled data in fam’ but not in nov (**Fig. 2H)**. These data jointly indicate that mPFC neurons largely retain their spatial tuning within the same environment but rapidly remap when experiencing a novel environment.

### A dorso-ventral gradient of spatial tuning

Hippocampal inputs are thought to provide a major source of spatially tuned afferents to the mPFC network (12). Tracing studies indicated that hippocampal projections to the mPFC are densest in ventral medial prefrontal areas (23) (24) (25) (26). Consistent with hippocampal projections to ventral mPFC areas, we found a higher power of theta (6-12 Hz) oscillations (Spearman’s r = -0.995, *P=*10^−63^, **Fig. 3A-C**) and a larger proportion of theta-coupled pyramidal cells in ventral mPFC areas (**SI Appendix Fig. S8**). We therefore hypothesized that spatial tuning of mPFC neurons located more ventrally in the mPFC might show stronger spatial tuning. Linear silicon probes allow pinpointing the location of the recorded units based on the amplitude of the extracellular waveform on neighboring electrodes, enabling us to determine the position of each recorded pyramidal neuron along the dorso-ventral axis (ranging from accessory motor cortex/cingulate cortex to prelimbic/medial orbital cortex, **Fig. 3A,D**). Unexpectedly, we detected stronger mean SI in the dorsal mPFC compared to ventral recording sites within individual mice (*N*=17 mice, *P*=0.03, paired *t-*test with Bonferroni correction for three comparisons; this adjustment was necessary as the data were also compared across two additional anatomical axes below; **Fig. 3D**). The larger SI values obtained from dorsal recording sites were not explained by a difference in the mean firing rates (28) (**SI Appendix Fig. S9**). We further observed a trend towards a larger proportion of pyramidal neurons with significant SI in the dorsal mPFC in individual mice (*P=*0.057, paired *t-*test with Bonferroni correction for three comparisons; **Fig. 3D, right**). The dorso-ventral SI gradient was also evident in the novel environment in individual animals (*P=*0.0064, **Fig. 3E**; as well as during re-exposure to fam’: *P=*0.036, *N=*11 mice, paired *t-*tests; **SI Appendix, Fig. S10**), indicating that it does not depend on prior experience with the track. Thus, spatial tuning emerges more prominently in the dorsal mPFC, which stays in marked contrast to the preferred projections of CA1 principal cells to the ventral mPFC (23) (24) (25) (26).

**Figure 3:**
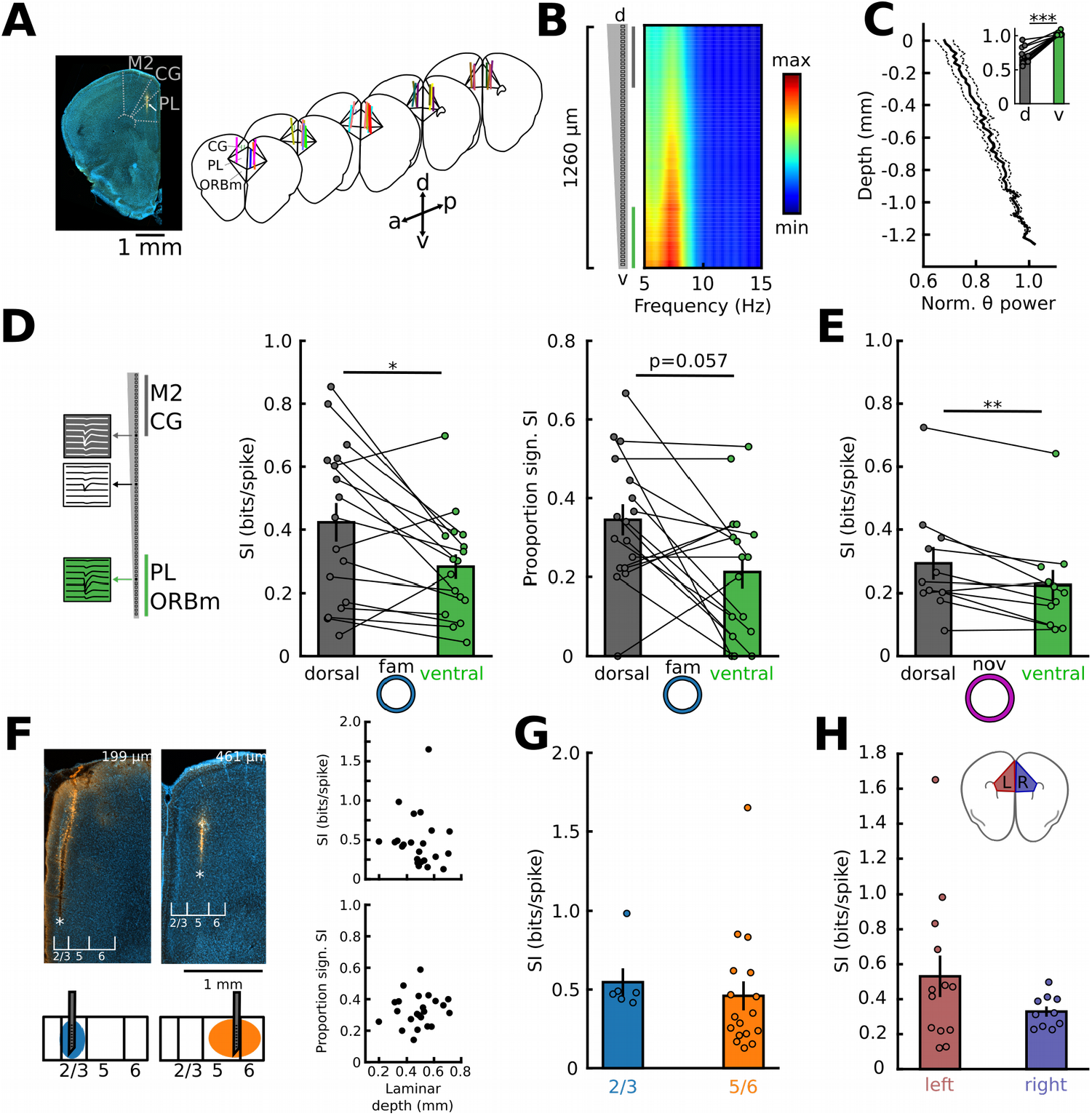
A dorso-ventral gradient in spatial representation. (A) Reconstruction of recording sites. Left: DiI-labelled probe track in the prefrontal cortex (arrow). Right: Location of silicon probes. Recording site for each mouse is presented in a different color (*N*=20 mice). M2: accessory motor cortex, PL: prelimbic cortex, CG: cingulate cortex, ORBm: medial orbital cortex, a: anterior, p: posterior, d: dorsal, v: ventral. (B) Example of the dorso-ventral depth profile of theta power along the shank. (C) Average normalized theta power as a function of d-v depth (*R=*-0.995, *P=*10^−63^). Inset: Average normalized theta power of the dorsal vs the ventral recording site (*P=*0.0004, Wilcoxon signed-rank test, *N=*16 mice). (D) Left: Identification of unit location along the probe shank. Grey and green depths indicate the locations used for grouping neurons into dorsal and ventral mPFC, respectively. Right: Comparisons of the mean SI (*P*=0.03) and the proportion of neurons with a significant SI for dorsal and ventral groups of cells recorded in individual mice (paired *t-*test with Bonferroni correction for 3 comparisons, *N=*17 mice). (E) The same comparisons as in (D) during exploration of the nov track (*P=*0.006, *N=*11 mice, paired *t-*test). (F) Comparable SI across cortical layers. Top: Histological examples of recording sites in superficial layers (2/3, left) and deep layers (5/6, right). Right: There was no correlation of the mean SI (*P*=0.256) or proportion of neurons with a significant SI (*P=*0.422) and laminar depth of the recording in individual mice (Spearman’s r, *N=*20 mice). (G) Comparable mean SI’s for superficial and deep layer recordings (*P=*0.076, Mann-Whitney U-test, *N*=6 and 17 mice). (H) The mean SI obtained from recordings in individual mice did not depend on the recording location in the left or right hemisphere (*P=*0.161, *N=*13 and 10 mice for left and right, respectively; unpaired *t-*test). Points in (C)-(H) represent average values for individual mice. Circles combined with lines represent data from individual mice. Continuous lines in (C) represent means across animals ± sem.

Previous work in the parietal cortex emphasized stronger spatial tuning in superficial layers (5). Since we so far analyzed recordings from both superficial (i.e., layers 2/3) and deep layers (i.e., layers 5/6), a laminar difference in SI might confound our observation of a dorso-ventral SI gradient. We performed a series of analysis to examine potential layer-specific effects: First, we compared SI values for shank locations in superficial and deep layers. The average SI and the proportion of neurons with significant SI per mouse did not depend on laminar depth (**Fig. 3F**, *P*=0.256 and *P=*0.422, *N=*20 mice, recording depth ranged from 199 to 669 µm). Furthermore, neither the mean SI nor the proportion of neurons with a significant SI differed between recordings conducted in layers 2/3 and 5/6 (SI: *P*=0.076, Mann-Whitney U-test; proportion sign. SI: *P=*0.777, unpaired *t-*test, *N*=6 and 20 mice for superficial and deep layers; **Fig. 3G; SI Appendix, Fig. S10**). Second, we performed simultaneous recordings at two cortical depths using multi-shank probes (shank spacing 250 µm). The mean SI did not depend on cortical depth in these recordings (**SI Appendix, Fig. S10B**). Finally, when layers were analyzed separately, a larger mean SI across cells was observed for dorsally located neurons for both superficial and deep recordings (**SI Appendix, Fig. S10C**). Jointly, these data argue against laminar differences in spatial tuning in the mPFC.

Previous investigations highlighted functional differences between the two hemispheres (29) (30) (31) (32) (33) (34) (35) (36). Motivated by our observed dorso-ventral gradient in spatial tuning, we therefore examined potential differences in SI between the left and the right hemisphere. We observed no difference in the mean SI or in the proportion of neurons with a significant SI recorded from the left and right hemisphere (*P=*0.161, unpaired *t-*test, *N=*13 and 10 mice for left and right hemisphere; **Fig. 3H, SI Appendix, Fig. S10D**). These data jointly demonstrate a gradient in the depth of spatial tuning along the dorso-ventral but neither laminar nor left-right axes.

We next asked whether the run-by-run reliability of spatial tuning might depend on the position along the dorso-ventral prefrontal axis. The average consistency of dorsally located cells was higher than that of ventral neurons in the fam environment (*P=*6.9*10^−5^, paired *t-*test, *N=*17 mice; **Fig. 4A,B**). Higher spatial consistency in the dorsal mPFC was also observed when mice were exposed to the nov environment (*P=*0.018, paired *t-*test, *N*=11 mice; **Fig. 4C**), suggesting that the higher reliability of spatial tuning of dorsal neurons did not depend on the context. In line with the more consistent spatial tuning of dorsal cells, decoding analysis allowed a more precise estimation of the animal’s position based on the firing of dorsal versus ventral neurons (*N=*5 mice, *P=*0.043, Wilcoxon signed rank test; **Fig. 4D**). Finally, we observed a trend towards higher mean stability in spatial tuning for dorsal vs. ventral neurons when we compared the first with the second session in the fam environment (fam vs fam’) for individual mice (*P*=0.056, paired *t-*test, *N=*11 mice, **SI Appendix, Fig. S11**), suggesting larger temporal stability of the spatial tuning properties in the dorsal mPFC. Taken together, our data jointly suggest that dorsally located mPFC neurons display more reliable and consistent spatial tuning, in particular within a given environment, than ventrally located cells.

**Figure 4:**
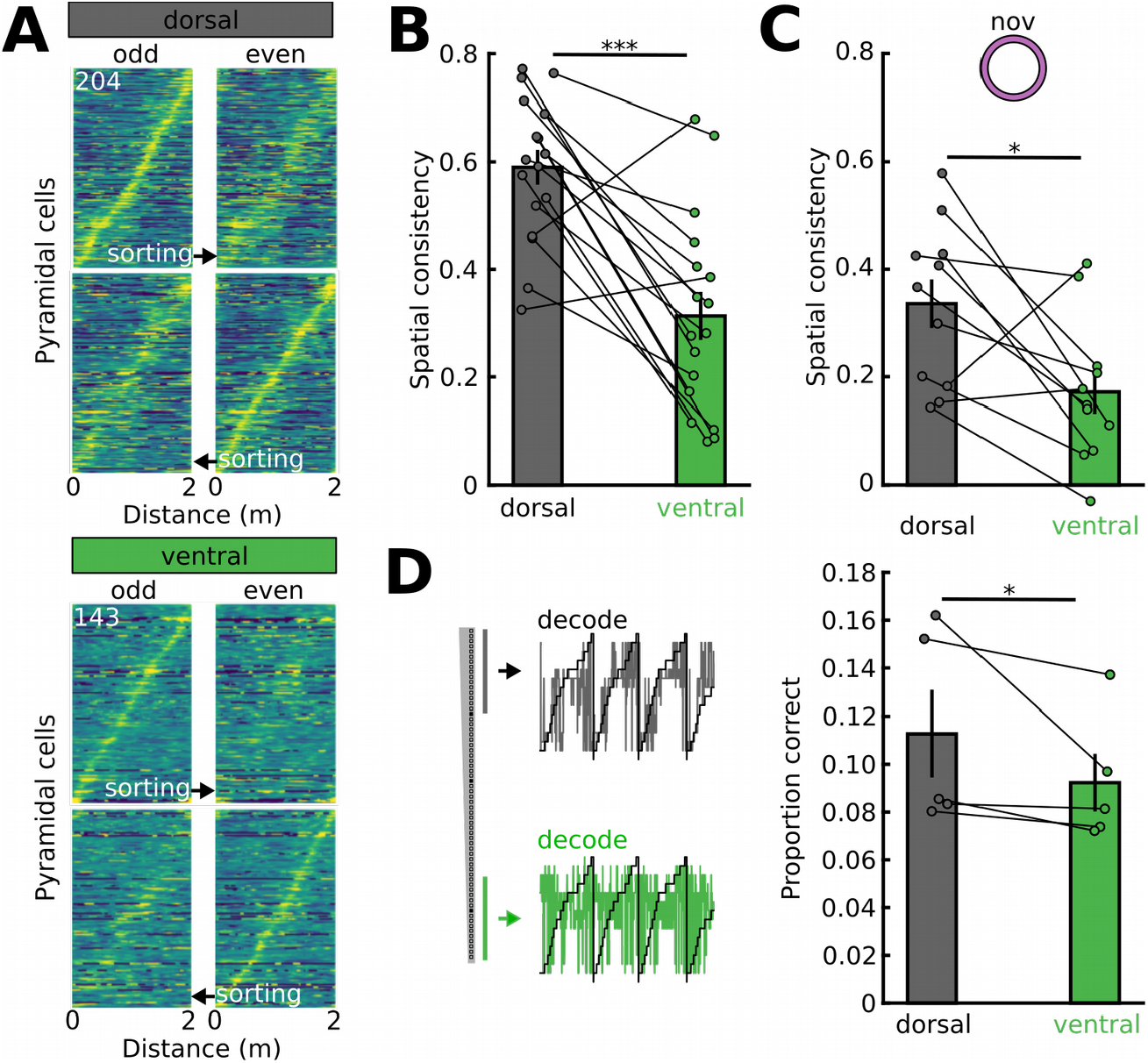
A dorso-ventral gradient in spatial consistency. (A) Tuning functions during alternating runs sorted for odd or even runs, separately for neurons belonging to the dorsal (top) our ventral group (bottom). (B) Higher spatial consistency (correlation between odd and even runs) of dorsally located neurons. *P=*6.9*10^−5^, paired *t-*test, *N=*17 mice. (C) Higher SI in dorsal neurons was also observed in the novel arena. *P=*0.018, paired *t-*test, *N=*11 mice. (D) Training a support vector classifier on dorsal neurons allowed a more precise prediction of the animal’s position than training the model on ventral neurons. *P=*0.043, Wilcoxon signed-rank test, *N=*5 mice. Circles in (B)-(D) show average values for individual mice.

## Discussion

Here we show that the mPFC of mice displays detectable spatial tuning during spontaneous, self-paced motion, including the representation of position as well as context. In line with findings of grid cells in the human cortex (37) (38) and spatially modulated cells in the cortex of rodents (4) (5) (6) (8) (9) (10), our findings are consistent with a more general neuronal system tracking spatial variables in neocortical columns (39).

We report that spatial tuning emerges in the mPFC in the absence of specific task rules or rewards. These results contrast some previous studies that found no apparent spatial tuning of prefrontal neurons in naturally exploring rodents (17) (19). Since prefrontal neurons modulate their firing rate in response to a variety of behavioural variables (e.g., grooming, rearing, (27)), it is possible that the reduced setting of the 1-dimensional circular track under head-fixation unmasked spatial tuning properties that were previously overlooked. The spatial tuning observed here might thus reflect a ‘default state’ of the network, which emerges when the mPFC is not engaged in particular task demands.

Analysis of trial-to-trial consistency revealed that the spatial tuning of individual mPFC neurons retains significant stability. This observation is supported by the ability to decode position based on spiking above chance level. However, it should be noted that the odd runs vs. even runs correlation of the tuning functions averages at ∼0.5. While similar consistencies were observed from our sample of hippocampal neurons, substantially higher consistency has been reported when the analysis is restricted to hippocampal place cells (20). Spatial tuning in the mPFC thus might be less pronounced and more transient in nature than in the hippocampus. Prefrontal consistency was larger when computed based on odd versus even runs (∼0.5) compared to runs during the first versus the second half of the recording (∼0.4, **SI Appendix, Fig. S6**), suggesting that in addition to variabilities in neuronal tuning during individual subsequent runs ‘representational drift’, i.e., slowly changing tuning of cortical neurons despite identical environment settings as observed in the parietal cortex (40), occurs in the mPFC.

It was previously unknown whether spatial tuning in the mPFC depends on prior experience. We have addressed that question by exposing mice to a novel track. We find similar SI compared to a familiar environment, which the mice have experienced for several consecutive days. Moreover, position decoding based on spike trains was equally effective in the novel environment. These data suggest that spatial tuning is established rapidly upon entering a novel environment, reminiscent of hippocampal CA1, in which place fields are observed immediately during novelty experience in the majority of neurons (41). In contrast, spatial tuning emerges gradually over the course of several days in the retrosplenial cortex (42), suggesting inter-area differences in the neocortex.

During novelty exposure, prefrontal spatial tuning displayed global remapping. These findings are again similar to results from hippocampal CA1, in which near-complete remapping is observed in novel environments in both freely moving (43) (44) and head-fixed mice (20). The observed neocortical remapping might thus be directly inherited from the remapping of hippocampal inputs. However, other mechanisms might also contribute. For instance, the mPFC is strongly innervated by dopaminergic afferents from the ventral tegmental area (VTA). Dopaminergic VTA neurons increase their firing rate in novel environments (45). While remapping in CA1 is not affected by stimulating dopaminergic VTA neurons (45), it remains to be investigated whether dopaminergic signals (or other neuromodulatory inputs) might impact prefrontal remapping.

The stronger spatial tuning of the dorsal mPFC is surprising since multiple studies demonstrated denser innervation of the ventral PFC by hippocampal inputs (23) (24) (25) (26), which presumably composes the primary source of spatial information to the mPFC (12). In addition to direct hippocampal afferents, spatially modulated inputs reach the mPFC via the thalamic nucleus reuniens, which relays hippocampal afferents to the mPFC (16) (46). However, similar to the direct hippocampal inputs, nucleus reuniens afferents have also been shown to preferentially target ventral prefrontal areas (46). Our findings of a dorso-ventral gradient of spatial tuning is thus not likely explained by direct hippocampal input. We propose that the spatial tuning of prefrontal neurons is only partly contingent on direct hippocampal input. Lesioning the hippocampus indeed reduced spatial information of neocortical cells and decreased position decoding accuracy based on spatially tuned neocortical neurons but did not entirely eliminate spatial tuning in the neocortex (10), consistent with an additional hippocampus-independent mechanism. Tracing studies confirmed the presence of entorhinal cortex projections in the dorsal and ventral portions of the mPFC (21) (22), which might contribute to spatial tuning properties of mPFC cells. In addition, spatial signals might reach the mPFC via other cortical areas. For instance, prominent spatial tuning features have been observed in the somatosensory (8) and orbitofrontal cortices (9), which project monosynaptically to the mPFC (47). Only few connections from the hippocampal formation to the somatosensory cortex have been identified (25), which gave rise to the hypothesis that the somatosensory cortex might contain a hippocampus-independent spatial navigation system (8). It remains to be tested whether the complex topographical nature of prefrontal spatial tuning emerges due to complementary inputs from the hippocampal and somatosensory spatial navigation systems.

Integration of spatial signals might be particularly relevant for movement-related tasks. Dorsal mPFC is strongly connected to downstream regions involved in sensorimotor functions. For instance, secondary motor and cingulate cortex (dorsal regions) of the rat project to the dorsal striatum, while medial orbital (also called infralimbic) cortex projects to ventral striatum, with prelimbic cortex taking an intermediate position with projections to both (48). Dorso-lateral striatum serves primarily sensorimotor functions, while ventro-medial striatum is most relevant for the integration of limbic signals (49). Along those lines, dorsal mPFC presumambly exerts a stronger direct influence on motor function than ventral areas as it contains a higher density of corticospinal neurons (48). Further investigations of area-specific mPFC circuits during cognitive tasks, resolving dorso-ventral subregions, will help to understand their contribution to cognitive behavior.

## Materials and Methods

### Mice

C57Bl6/J mice of both sexes were maintained on a 12h dark-light cycle with free access to food and water. At the start of the experiment, the animals were 6 to 13 weeks old. All experiments were performed in agreement with national legislation and were approved by the Regierungspräsidium Freiburg.

### Surgical procedures

For single-unit recording in the virtual reality, a stainless steel head plate was implanted on the skull under general anesthesia in isoflurane (induction: 3%, maintenance: 1-2%) using dental cement. The animals were allowed to recover from head plate implantation for at least three days. Buprenorphin (0.1 mg/kg body weight) and Carprofen (5 mg/kg body weight) were injected subcutaneously before the surgery for pain relief. Once the animals were habituated to head-fixation (see below), a craniotomy was performed over both mPFCs (1.9 mm anterior, 0.4 mm lateral of bregma) under isoflurane anesthesia. The craniotomy was then sealed of with QuikCast elastomer until the recordings took place. An additional injection of Carprofen was given for analgesia prior to craniotomy.

### Single-unit recording in the virtual reality

After recovery from head-plate implantation, the mice were trained to run on a circular track in a virtual reality. They were briefly sedated with isoflurane applied in oxygen and head-fixed such that they could comfortably stand on a circular styrofoam weel. First, animals were habituated to head-fixation without the virtual reality turned on. After at least three days of habituation, training in the virtual reality (circular track, length 1.5 m or 2 m, visual cues placed outside the arena) was conducted. The virtual reality was constructed with open-source 3D rendering software Blender (50) and was projected on five computer screens surrounding the head-fixation setup. Over subsequent days, mice were accustomed to the virtual environment until the animals appeared calm and traversed the circular maze reliably.

Recordings were performed in the arena to which the animals were habituated (*N*=20 mice mPFC, *N*=2 mice CA1). 3 of the mice with mPFC recording had an additional local field potential wire implanted into the olfactory epithelium as part of a different study. The animals completed on average 28.1 ± 5.9 laps per recording session. In a subset of mice (*N*=11 animals), we additionally exposed the animals within the same session to a novel arena (nov, 14.3 ± 2.0 laps completed), which the mice encountered for the first time during the initial recording session. The novel arena differed in size (3 m circle length) and visual features from the familiar arena. After recording in the novel arena, the animals were again exposed to the familiar track to assess the reactivation of spatially tuned activities in the familiar context (25.4 ± 6.3 laps were completed on average, *N*=11 mice).

Recordings were conducted 1-3 days after the craniotomy. A single-shank silicon probe with 64 recording sites (electrode pitch 20 µm, total shank length 1275 µm, H3 probe, Cambridge Neurotech) coated with a fluorescent marker (DiI or DiO) was slowly (∼2-5 µm/s) lowered to the mPFC (1800-1900 µm below brain surface). In three mice, a four shank probe with 16 electrodes per shank (Cambridge Neurotech) was used to perform simultaneous recordings from different neocortical layers. After insertion, the probe was left in place before starting the recording for 10-15 min. In two mice, recordings were obtained from hippocampal CA1. Wide-band neural signals were recorded at 30 kHz sampling with a 64-channel amplifier (Intan Technologies) connected to an USB acquisition board (OpenEphys). The position in the virtual reality was recorded as a pulse-width modulated signal using one of the analog input channels of the OpenEphys USB interface board. After recording, the silicon probe was slowly retracted and the craniotomy sealed off with QuickCast elastomer. 1-2 recording sessions were performed with each animal, with one session per day.

### Histology

After the last recording session, the animals were deeply anesthetized with an intraperitoneal injection of ketamin/xylazine and transcardially perfused with ∼20 ml phosphate-buffered saline followed by ∼30 ml of 4% paraformaldehyde. After post-fixation over night, 100 µm-thick frontal sections of the mPFC were cut, washed several times in phosphate-buffered saline (PBS) and stained with 4′,6-diamidino-2-phenylindole. The location of the silicon probe was visualized with a laser-scanning microscope (LSM 710 or 900, Zeiss) following the nomenclature of the Allen Brain Atlas. To determine laminar depth of the probe, the distance from pia to the shank was measured orthogonal to the cortical layers.

### Single-unit isolation

Single unit clusters were isolated from bandpass-filtered raw data (0.3-6 kHz) using MountainSort (51). Putative single-unit clusters that fulfilled quality criteria of high isolation index (>0.90) and low noise overlap (<0.1) were kept for manual curation, during which only clusters with a clear refractory period and clean waveform shape were saved for further analysis. In case of two clusters with similar waveforms, cross-correlation was used to assess whether clusters had to be merged. Isolated units were separated in putative excitatory and inhibitory neurons based on trough-to-peak duration and asymmetry index (52) using k-means clustering. To quantify the stability of single-unit recording during the recording sessions, waveforms of the channel with the largest voltage deflection were average separately for the time the animal explored the familiar track and during re-exposure to the familiar track after completion of running on the novel track. Stability was assessed by calculating Pearson’s correlation coefficient between the waveforms of each unit during both conditions and by correlating the peak amplitude recorded during both sessions. To extract the physical location of the units, the channel on the silicon probe with the largest negative amplitude deflection was defined as the location of the unit and matched to histological data. All spike analysis was performed during times when the animal was moving at a speed of at least 2 cm/s.

### Analysis of single unit data

To minimize the impact of low firing rate on SI, only neurons with an average firing rate above 1 Hz were considered for analysis. Spike data was first aligned to the position signal in the maze. For that, the pulse-width modulated signal indicating position was transformed to a continuous signal ranging from 0 to 1 (denoting start and end of a complete circle) by low-pass filtering and manual curation of start and end points. Firing rate during movement on the maze was estimated by taking all spikes during which the minimum speed criterion was fulfilled and dividing them by the total time spent moving. The spiking of individual units was binned as a function of space on the circular maze (2 cm bin width) and normalized by occupancy of each bin. The resulting average tuning function of each cell was smoothed with a Gaussian filter of SD=4 cm. The neurons were sorted by the peak location in the smoothed spatial tuning function for display in **Fig. 1**. SI was calculated as

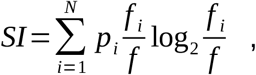

where *N* is the number of bins, *p* is the occupation probability in the i^th^ bin, *f*_*i*_ is the activity in bin *i*, and f is the average activity of the neuron across bins (10) (28). A control data set was constructed for each neuron by randomly shifting each spike between 10 and 30 s. Time-shifts of 5-20 s or 30-40 s as well as different spatial binning (4 cm with 8 cm spatial smoothing) gave similar results (**SI Appendix, Fig. S5**). The proportion of neurons with significant SI was obtained by calculating SI for shuffled spike train 1000 times. A cell was scored as expressing significant SI when the measured SI exceeded the 99^th^ percentile of the shuffled distribution. All further analysis, including decoding, was applied to all neurons, irrespective of whether or not they expressed significant SI. To determine spatial consistency within the session, average occupancy-normalized spatial histograms of spikes were constructed separately for odd and even runs of a given recording session. Consistency was then defined by Pearson’s correlation coefficient between both histograms. Alternatively, spatial histograms of spikes were constructed separately for the first and second half of the recording. For spatial stability, average spike rates as a function of position on the maze were obtained separately for fam, nov and fam’. Stability was defined as Pearson’s *r* computed for the average spatial maps.

Theta coupling was assessed by first extracting the phase of the 6-12 Hz-filtered local field potential recorded in the middle of the shank during running, selecting cells with non-uniform phase distributions using Rayleigh’s test, and computing the pairwise phase consistency *PPC (53)* as

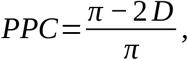

where *D* is the average pairwise circular distance between individual spikes as

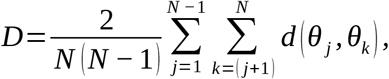

where *θ*_*j*_ and *θ*_*k*_ are the phases of the local field potential of spikes *j* and *k*, and *N* is the total number of spikes of the unit. Theta power was computed using Welch’s method with a window size of 10 s and 10x zero-padding.

To assess whether prefrontal neurons are speed- and/or acceleration-modulated, spike trains of neurons were binned as a function of instantaneous speed/acceleration obtained from position data (30 bins). Modulation was then assessed by correlation between the speed/acceleration-binned spike trains and the mean speed/acceleration of each bin (Spearman’s r).

### Decoding analysis

Position decoding was performed with a linear support vector classifier using the scikit learn package (sklearn.svm.LinearSVC with C=5). First, the instantaneous firing rate of each pyramidal cell was obtained by convolving the spike train with a Gaussian kernel of SD=50 ms. The data were then downsampled by a factor of 500 to reduce computation time. The position signal (normalized from 0 to 1) was digitized in 10 cm bins. We then performed an iterative procedure in which the instantaneous rate of a random number of pyramidal units was selected from the total population 100 times (ranging from 2-20 neurons per iteration). To decode within an environment, decoding accuracy was expressed as the mean model accuracy obtained with 5-fold cross-validation averaged over all iterations with the same number of neurons. Note that this is a very conservative measure of performance, since for each time point only the prediction of the single correct spatial position bin is scored as correct. Control data to estimate chance level was generated for each recording session by randomly shuffling the position signal. For visualization of decoding and for the estimation of maximal decoding performance in **Fig 1G** and **Fig. 2G**, only session with >30 simultaneously recorded pyramidal cells were considered (56 ± 4 neurons in fam, *N=*11 mice, 57 ± 7 neurons in nov, *N=*8 mice). To decode across environments, the spiking and discretized position data of each environment (20 bins) were concatenated, and decoding was tested using sklearn’s GroupKFold data splitting such that training and testing data came from the two distinct environments.

### Statistical analysis

Unpaired comparisons were performed with a Mann-Whitney U-test if the data did not follow a normal distribution, otherwise an unpaired *t*-test was used. Normality was assessed with a Shapiro-Wilk test. Pairwise comparisons were done with a paired *t-*test or Wilcoxon signed-rank test (when data did not follow a normal distribution). Bonferroni correction was applied for multiple comparisons (e.g. by correcting the critical *P=*value from 0.05 to 0.016 when comparing SI across three anatomical axis). Correlations between spatial tuning functions were tested with Pearson’s correlation coefficient to detect linear correlations. Correlations for which linearity could not be assumed (e.g. cortical depth *vs*. spatial information) were assessed with Spearman’s correlation coefficient. Significant theta coupling was assessed with Rayleigh’s test for circular uniformity. Data are presented as mean ± sem. All analysis (except for initial spike sorting, see above) including statistics were performed in Python (versions 2.7 and 3.7).

## Acknowledgments

We thank Johannes Letzkus, Jozsef Csicsvari and Antje Kilias for critical comments on earlier versions of the manuscript, and Kerstin Semmler and Karin Winterhalter for technical assistance.

