## Supplemental Information for "Topographically organized representation of space and context in the medial prefrontal cortex"

Supplemental Figures 1-11

Supplemetal Table 1

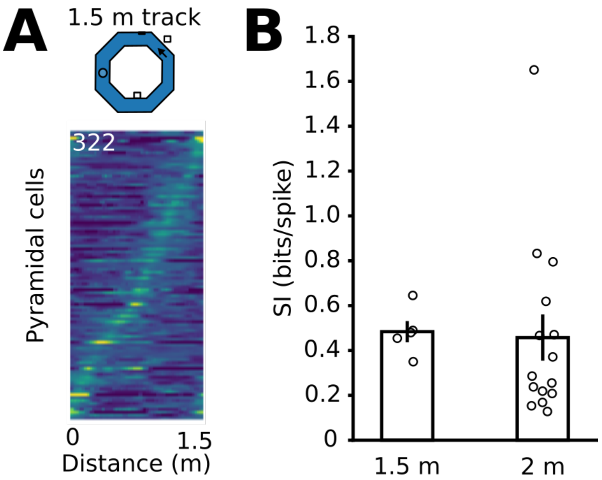


**Fig. S1: Spatial tuning emerges across scale of the familiar environment.**

(A) Peak sorted spatially binned firing activity of pyramidal cells during recordings on a shorter circular track (1.5 m long, $N$=5 mice).
(B) SI did not differ between short and long familiar tracks. $N$=5 and 15 mice, $P$=0.128, Mann-Whitney U-test. Each dot denotes average SI of one animal. Bars show the mean ± sem.

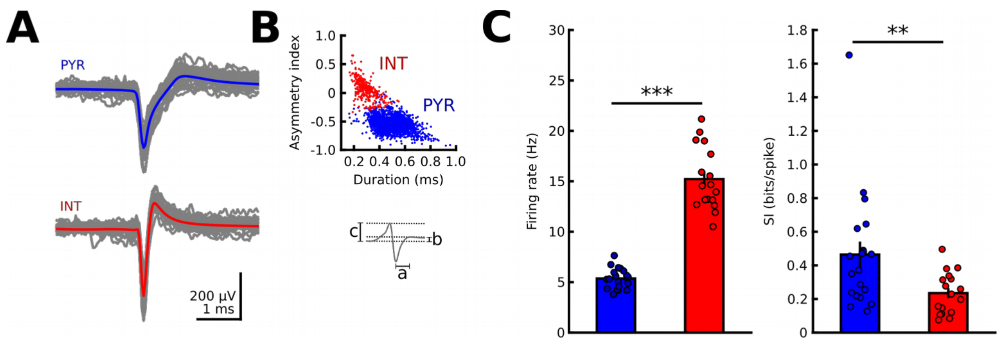


**Fig. S2: Identification and spatial tuning of prefrontal interneurons.**

(A) Example waveforms of a pyramidal cell (PYR, top) and a putative interneuron (INT, bottom). Grey traces show 30 randomly selected single waveforms, blue and red traces are averages.
(B) Classification of PYR and INT based on duration *a* and asymmetry index (*b*-*c*)/(*b*+*c*) of filtered action potential waveforms. Each dot shows a single unit. *N*=2005 PYR and 324 INT.
(C) Putative INT showed significantly higher firing rates than PYRs (*N*=20 and 17 mice, $P=10^{-15}$, Welch's test). Right, INT show significantly lower SI than PYR (N=20 and 17 mice, *P*=0.005, Mann-Whitney U-test). Each denotes average values of a single animal. Bars show the mean ± sem.

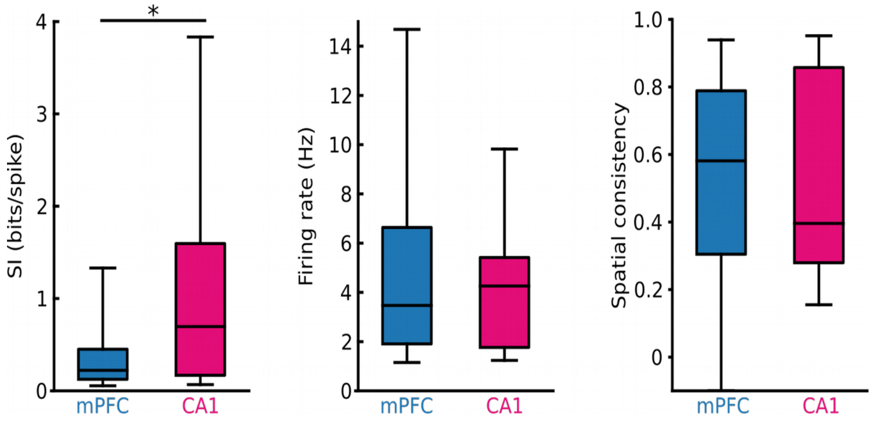


**Fig. S3: Comparison of spatial tuning between prefrontal and hippocampal pyramidal cells.**

SI of prefrontal pyramidal neurons was significantly lower than in CA1 (left, *P*=0.0012), while no difference was found for firing rate (middle, *P*=0.481) or consistency in spatial tuning between odd and even runs (right, *P*=0.457). *N*=1272 and 26 cells from 20 and 2 mice, respectively; Mann-Whitney U-tests. * *P*<0.05.

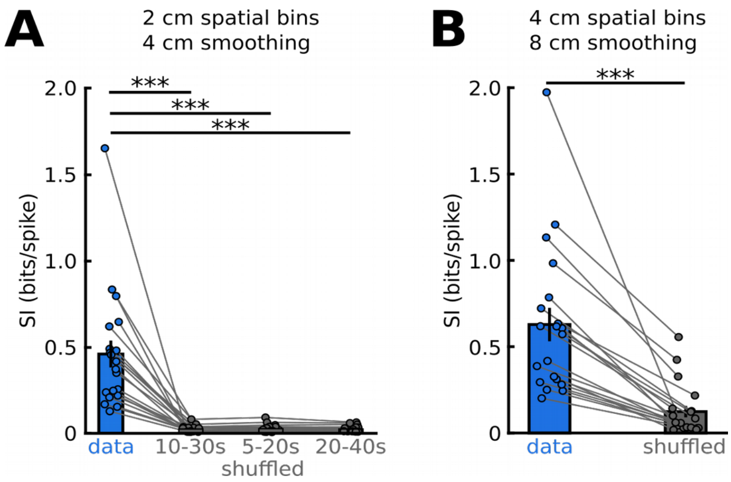


**Fig. S4: Consistent SI for different shuffling and spatial binning parameters.**

(A) Average SI for prefrontal neurons compared to various spike time-shift controls. Each spike was independently shifted forward in time between 10-30, 5-20 or 20-40 s in the three shuffled control groups. $P$=4.8*$10^{-5}$, 5.1*$10^{-5}$ and 4.8*$10^{-5}$, paired $t$-test with Bonferroni correction for three comparisons. $N$=20 mice

(B) Significant SI did not depend on the spatial binning as it was also observed for wider 4 cm bins. $P$=1.0*$10^{-5}$, $N$=20 mice. Each dot in (A) and (B) shows the average SI value for an individual mouse. Circles combined by lines were obtained within one animal. Bars show the mean ± sem.

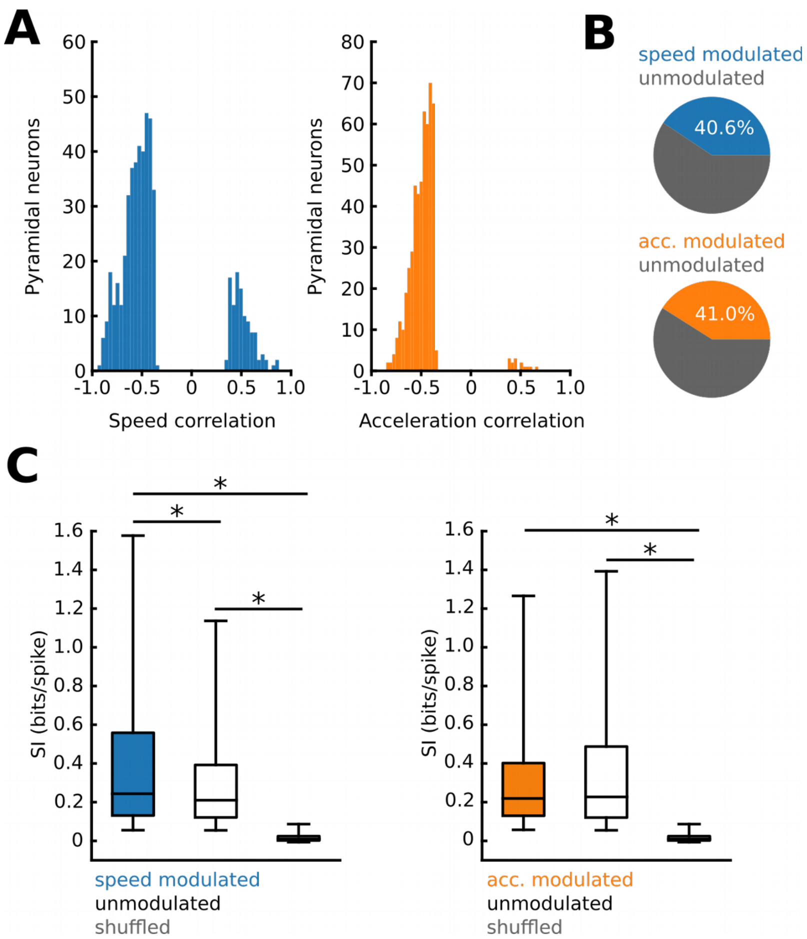


**Fig. S5: Speed modulation of prefrontal pyramidal cells.**

(A) Histogram of speed (left) and acceleration (right) correlation values of all pyramidal cells. Only neurons with significant modulation determined by Spearman's rank correlation coefficient are shown.
(B) ~40 % of prefrontal neurons were either modulated by speed or acceleration.
(C)) Significant SI of prefrontal neurons persists when speed- or acceleration-modulated cells are removed from the calculation (white boxes, compared to shuffled controls shown in grey).

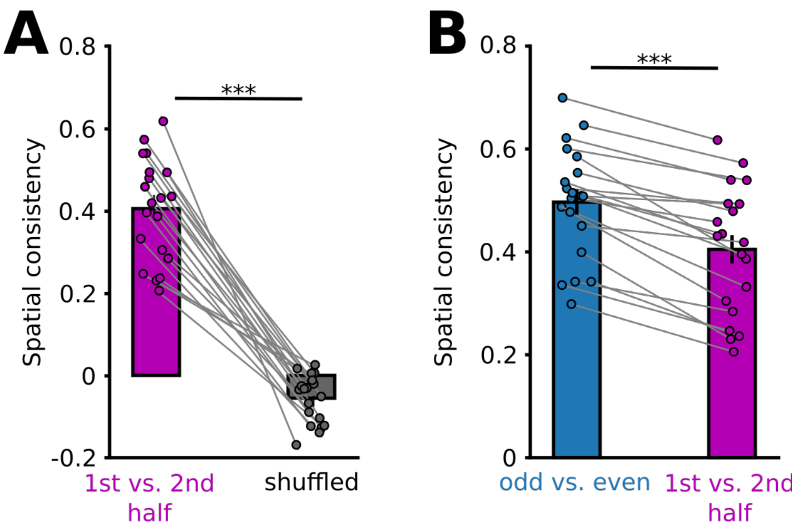


**Fig. S6: Significant spatial consistency based on the first and the second half of runs.**

(A) Spatial consistency was computed using the first and second half of runs, which revealed significant consistency compared to shuffled controls. T=13.897, $p=2*10^{-11}$, paired t-test, N=20 mice. Each point shows the average consistency of all pyramidal neurons of one mouse.

(B) Consistency was higher using the odd versus even runs compared to the first half versus second half method, suggesting a temporal drift in spatial tuning of prefrontal pyramidal neurons. T=8.218, $p=1*10^{-7}$, paired t-test, N=20 mice.

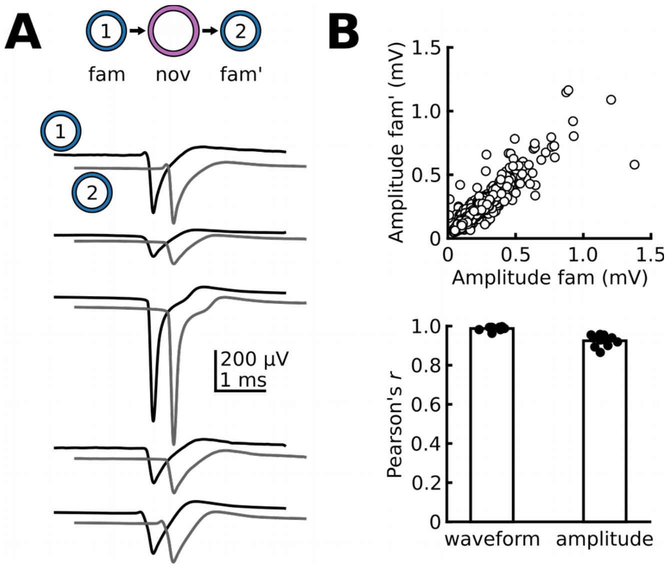


**Fig. S7: Additional analyses of waveform stability during exposure to nov and fam'.**

(A) To assess recording stability, the average waveform of each unit was calculated during exploration of the familiar track (fam) and during re-exposure to the familiar track (fam'). Bottom: Examples of average waveforms of five units.
(B) Top: Peak amplitudes of all units during fam and fam' were highly correlated. *N*=1493 units, 11 mice, Pearson's $r$ = 0.912. Bottom: Summary of waveform correlation and peak amplitude correlation during each session. *N*=11 mice). Bars show the mean and sem.

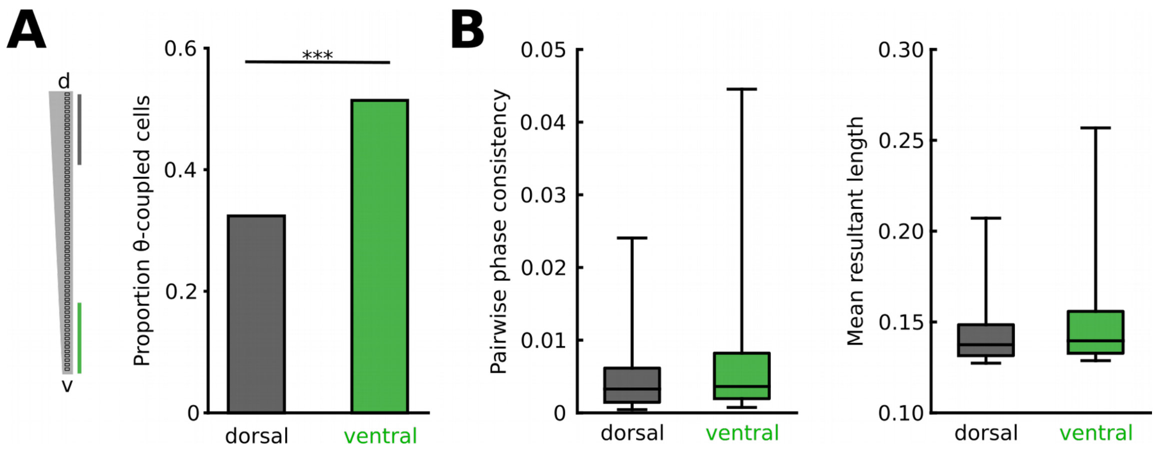


**Fig. S8: Theta-coupling along the dorso-ventral axis.**

(A) The proportion of theta coupled neurons was significantly larger in dorsal than ventral mPFC areas. Dorsal: $N$=88 of 272 neurons coupled, ventral: 113 of 220 neurons coupled, $P=2.2*10^{-5}$, Fisher's exact test. Neurons with significant SI did not differ between superficial and deep layers.
(B) Coupling depth quantified as pairwise phase consistency (left, $P$=0.606) or mean resultant length (right, $P$=0.415) did not differ for dorsal and ventral groups. $N$=88 dorsal and 113 ventral neurons, Welch's tests (17 mice).

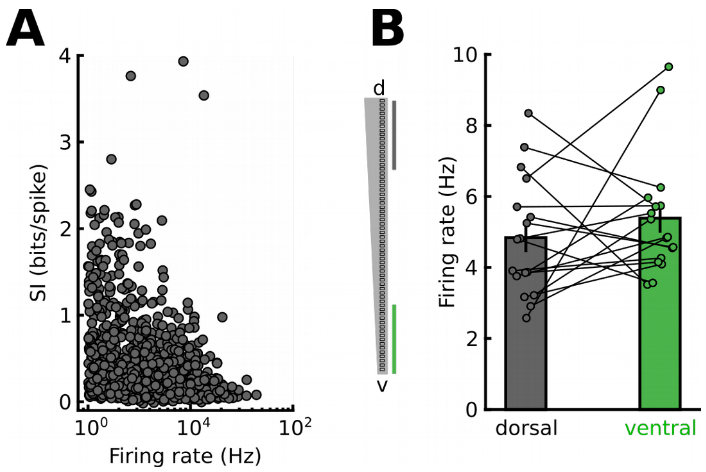


**Fig. S9: Firing rate dependence of SI.**

(A) Consistent with previous reports, SI was inversely correlated with firing rate. Spearman's r = -0.34, *P*=2*10$^{-35}$. Each circle denotes a pyramidal cell.
(B) Average firing rate did not differ between dorsal (grey) and ventral (green) recording sites, arguing against firing rate differences as a confounding factor towards larger SI in dorsal areas. Data points represent averages of individual mice (*P*=0.337, *N*=17 mice, paired *t*-test). Circles combined by lines were obtained within one animal.

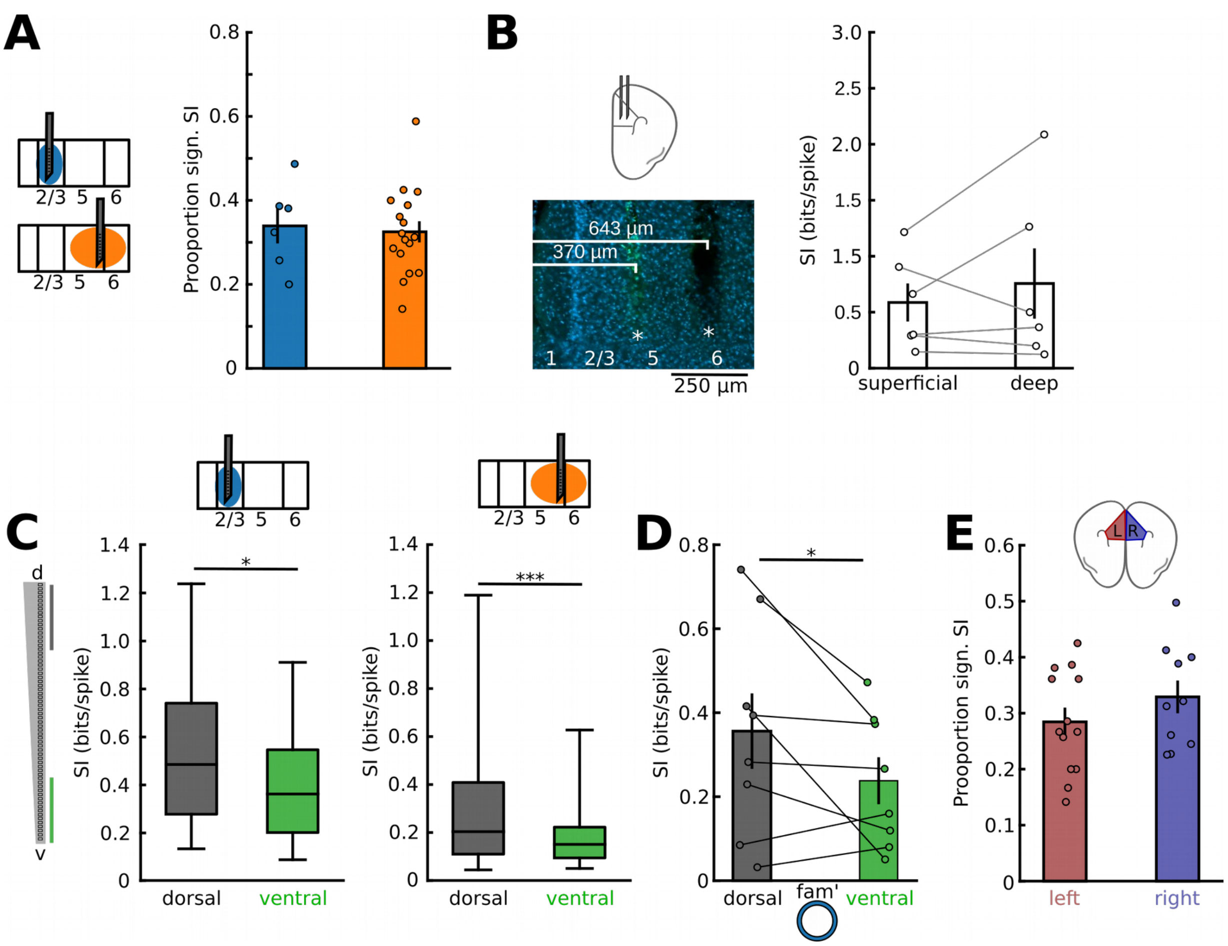


**Fig. S10: Similar spatial tuning across the superficial-deep and left-right axes.**

(A) The proportion of neurons with a significant SI did not differ between superficial (*N*=6 mice) and deep layers (*N*=20 mice) (unpaired *t*-test, *P*=0.777).
(B) Top left, schematic of simultaneous recordings with two shanks placed at different cortical depths. Bottom left, example of a dual-site recording at 370 and 643 µm depth. Right, summary of average SI's of neurons recorded on superficial and deep shanks. Values are session averages for both shanks from *N*=3 mice (*P*=0.422, paired *t*-test, *N*=6 sessions). Bars represent the mean ± sem.
(C) Box plot summarizes SI for superficial (left, *N*=67 and 69 neurons) and deep layer neuron recordings (right, *N*=205 and 148 neurons) and reveals significantly larger SI in dorsal cells (*P*=0.019 and $P=1.7*10^{-5}$, Welch's tests).
(D) Significantly larger SI for dorsal neurons during exploration of fam'. *N*=11 mice, *P*=0.036, paired *t*-test).
(E) The mean proportion of neurons with a significant SI in individual mice did not differ between the left and the right hemisphere (*N*=13 and 10 mice, *P*=0.161, unpaired *t*-test). Circles in (A), (B) and (D) are averages of individual mice. Circles combined by lines were obtained within one animal. Boxplots in (C) are based on cells. **P*<0.05, ****P*<0.001.

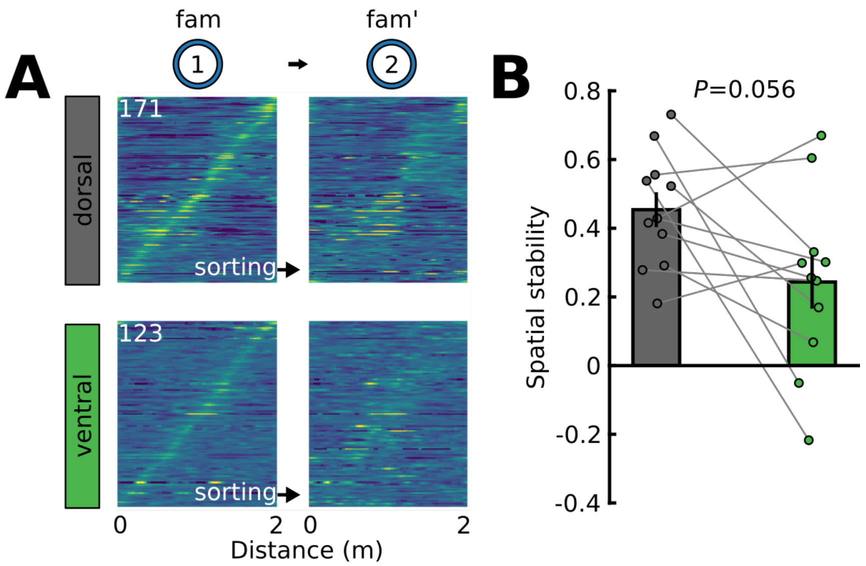


**Fig. S11: Stability of spatial tuning across fam-fam’ for dorsal and ventral prefrontal neurons.**

(A) Peak-sorted pyramidal cell spatial tuning functions during exploration of two subsequent familiar environments (sequence of runs: fam, nov, fam’; sorting was kept the same), separately for dorsal (top, grey) and ventral (bottom, green) prefrontal neurons. Runs within the nov environment positioned between fam and fam’ are not shown for clarity.
(B) There is a trend towards higher mean spatial stability (fam vs fam’) for dorsal neurons, which did not reach significance (*N*=11 mice, paired *t*-test). Circles represent average values of individual mice. Circles combined by lines were obtained within one animal.

**Table S1: Summary of values and statistics**

| Data | Mean ± sem | subjects | test | Statistic | *P* | comment |
|---|---|---|---|---|---|---|
| Spatial tuning in the familiar environment (Fig. 1) | | | | | | |
| SI | 0.46 ± 0.08 | 20 mice | Paired t-test | t=5.825 | $1.3*10^{-5}$ | |
| shuffled | 0.02 ± 0.004 | | | | | |
| consistency | 0.497 ± 0.025 | 20 mice | Paired t-test | t=19.22 | $6.5*10^{-14}$ | |
| shuffled | -0.061 ± 0.013 | | | | | |
| decoding accuracy | 0.276 ± 0.022 | 11 mice | Paired t-test | t=9.97 | $1.6*10^{-6}$ | 56 ± 4 cells |
| shuffled | 0.075 ± 0.006 | | | | | |
| Remapping in the novel environment (Fig. 2) | | | | | | |
| stability fam-nov | 0.057 ± 0.028 | 11 mice | Paired t-tests | 8.363 | $2.4*10^{-5}$ | Bonferroni correction for 3 comparisons |
| stability fam-fam’ | 0.362 ± 0.045 | | | | | |
| shuffled | 0.075 ± 0.029 | | | 4.595 | 0.003 | |
| decoding in nov | 0.147 ± 0.022 | 8 mice | Paired t-test | t=0.514 | 0.614 | 57 ± 7 cells |
| shuffled | 0.041 ± 0.002 | | | | | |
| decoding fam-nov (fold chance) | 1.533 ± 0.084 | 11 mice | Paired t-test | t=1.149 | 0.277 | 20 cells vs chance |
| decoding fam-fam’ (fold chance) | 1.055 ± 0.046 | 11 mice | Paired t-test | t=5.847 | 0.002 | 20 cells vs chance |
| Dorso-ventral SI gradient (Fig. 3) | | | | | | |
| Theta power dorsal | 0.687 ± 0.029 | 16 mice | Paired t-test | t=0 | 0.0004 | Normed to ventral |
| Theta power ventral | 1.020 ± 0.005 | | | | | |
| D-v correlation in theta power | | 16 mice | Spearman’s r | r=-0.995 | $10^{-63}$ | |
| SI dorsal | 0.423 ± 0.061 | 17 mice | Paired t-test | t=2.915 | 0.03 | Bonferroni correction for 3 comparisons |
| SI ventral | 0.282 ± 0.039 | | | | | |
| Prop. sig. SI dorsal | 0.345 ± 0.040 | 17 mice | Paired t-test | t=2.519 | 0.057 | Bonferroni correction for 3 comparisons |
| Prop. sig. SI ventral | 0.212 ± 0.040 | | | | | |
| SI dorsal (nov) | 0.293 ± 0.052 | 11 mice | Paired t-test | t=3.432 | 0.006 | |
| SI ventral (nov) | 0.225 ± 0.047 | | | | | |
| SI dorsal (fam’) | 0.324 ± 0.067 | 11 mice | Paired t-test | t=2.411 | 0.036 | |
| SI ventral (fam’) | 0.213 ± 0.042 | | | | | |
| Correlation SI – laminar depth | | 20 mice | Spearman’s r | | 0.256 | |
| Correlation prop. sig SI – laminar depth | | 20 mice | Spearman’s r | | 0.422 | |
| SI superficial | 0.547 ± 0.088 | 6 mice | Mann- | U=30 | 0.076 | |

| | | | | | | |
|---|---|---|---|---|---|---|
| SI deep | 0.460 ± 0.092 | 17 mice | Whitney U-test | | | |
| Prop. sig. SI superficial | 0.339 ± 0.042 | 6 mice | Unpaired t-test | t=0.287 | 0.777 | |
| Prop. sig. SI deep | 0.325 ± 0.025 | 17 mice | | | | |
| SI left | 0.531 ± 0.119 | 13 mice | Unpaired t-test | t=1.454 | 0.161 | |
| Si right | 0.314 ± 0.052 | 10 mice | | | | |
| Dorso-ventral consistency gradient (Fig. 4) | | | | | | |
| Consistency dorsal | 0.590 ± 0.032 | 17 mice | Paired t-test | t=5.320 | $6.9*10^{-5}$ | |
| Consistency ventral | 0.313 ± 0.045 | | | | | |
| Consistency dorsal (nov) | 0.336 ± 0.045 | 11 mice | Paired t-test | t=2.817 | 0.018 | |
| Consistency ventral (nov) | 0.172 ± 0.040 | | | | | |
| Decoding accuracy dorsal | 0.113 ± 0.018 | 5 mice | Wilcoxon signed-rank test | t=0 | 0.043 | 5 dorsal and ventral cells used for each mouse |
| Decoding accuracy ventral | 0.092 ± 0.012 | | | | | |
